# Colorectal Cancer-Associated *Streptococcus gallolyticus*: A Hidden Diversity Exposed

**DOI:** 10.1101/2025.02.28.640884

**Authors:** Bruno Périchon, Thomas Cokelaer, Wooi Keong Teh, Laurence du Merle, Laurence Ma, Alexandra Doloy, Claire Poyart, Michael Givskov, Patrick Trieu-Cuot, Shaynoor Dramsi

**Affiliations:** Institut Pasteur, Université Paris Cité, Unité de Biologie des Bactéries Pathogènes à Gram-Positif, Paris Cedex 15, France; Institut Pasteur, Université Paris Cité, INSERM U1201, Unité de Parasitologie moléculaire et Signalisation, F-75015 Paris, France; Institut Pasteur, Université Paris Cité, Bioinformatics and Biostatistics Hub, F-75015 Paris, France; Institut Pasteur, Plateforme technologique Biomics, F-75015 Paris, France; Assistance Publique Hôpitaux de Paris, Hôpitaux universitaires de Paris Centre-site Cochin Service de Bacteriologie, Centre National de Référence des Streptocoques, F-75014 Paris; Singapore Centre for Environmental Life Sciences Engineering, Nanyang Technological University, Singapore, Singapore; Costerton Biofilm Center, Departement of Immunology and Microbiology, Faculty of Health and Medical Sciences, University of Copenhagen, Denmark; Université Paris Cité, CNRS UMR8104, Inserm U1016, Institut Cochin, Bacterial Pathogenesis and Innate Immune Signaling Team, F-75014 Paris, France

**Keywords:** *Streptococcus gallolyticus* subsp. *gallolyticus*, genome, antibiotic resistance genes, *Streptococcus gallolyticus* subsp. *macedonicus*, colorectal cancer

## Abstract

*Streptococcus gallolyticus* subsp. *gallolyticus* (*SGG*) is a bacterial pathogen implicated in bacteremia and endocarditis, and is often associated with colon tumors in elderly individuals. The development of colorectal cancer (CRC) has been linked to intestinal dysbiosis, characterized by increased proportions of *SGG* and other intestinal microbes. In this study, we present the complete nucleotide sequence of five novel clinical isolates of *SGG* associated with colorectal cancer, revealing unexpected genetic diversity. Sequencing an additional 30 *SGG* clinical isolates provided a more comprehensive description of this genetic diversity. We did not identify a pathogenicity island specific to CRC-associated *SGG* isolates. Most of these human-derived *SGG* isolates exhibit resistance to multiple antibiotics. Our findings also offer additional insights into multilocus sequence typing (MLST), capsular loci, and pilus organization. Analysis of the repertoire of surface proteins reveals high potential for binding and foraging complex polysaccharides. Finally, comparative genomics with the phylogenetically closest non-pathogenic subspecies *S. gallolyticus* subps. *macedonicus,* confirmed that *SGG* pathogenicity-associated factors mostly rely on a large repertoire of surface proteins involved in host colonization, presence of C5a peptidase to avoid innate immunity, bile salt hydrolase to persist in the gut, and of specific bacteriocin and type VII-dependent effectors to colonize the host colon. Additionally, the presence of extracellular polysaccharides in *SGG* probably helps the bacterium survive in harsher conditions.

**Importance:** *Streptococcus gallolyticus* subsp. *gallolyticus* (*SGG*) was the first intestinal bacterium associated with colorectal cancer. It is now widely accepted that colonic microbiota dysbiosis contributes to oncogenesis, with a higher relative abundance of several potentially pro-carcinogenic bacteria. For example, the oncogenic role of *Escherichia coli* pks+ and enterotoxinogenic *Bacteroides fragilis* in colorectal cancer have been well established identifying the role of genetic loci encoding toxins.

Through the sequencing and analysis of 11 clinical *SGG* isolates from CRC patients and comparisons with non-CRC isolates, we uncovered a significant diversity among CRC-associated strains. Our findings suggest that *SGG* association with CRC is complex and is not linked to a specific strain or pathogenicity island, thus highlighting the opportunistic and versatile nature of *SGG*.

## Introduction

*Streptococcus gallolyticus* subsp. *gallolyticus* (*SGG*), formerly known as *Streptocococcus bovis* biotype I, is a member of the larger *Streptococcus bovis*/*Streptococcus equinus* complex (SBSEC). The group *Streptococcus gallolyticus* includes two additional subspecies, namely subsp. *macedonicus* (*SGM*) and subsp. *pasteurianus* (*SGP*).

*SGG* is a commensal bacterium of the digestive tract of birds and of the rumen of herbivores (1). It is estimated that around 30% of humans harbor this bacterium in their gastrointestinal tract (2). *SGG* is recognized as an opportunistic pathogen, primarily responsible for bacteremia and endocarditis, especially in elderly individuals.

The first whole genome sequence of *S. gallolyticus* subsp. *gallolyticus* strain UCN34 was published in 2010 (3). This clinical isolate was recovered from the blood of a 70-year-old man with endocarditis and a colorectal tumor. Genome analysis of strain UCN34 revealed distinct features and suggested that this subspecies likely acquired genes through horizontal gene transfer from other species present in the rumen flora, such as *Lactobacillus* or *Clostridia* species (3).

*SGM*, a species used in the dairy industry, is the genetically closest relative to *SGG* and is considered non-pathogenic. Comparative genomics of *SGG* and *SGM* allowed the identification of putative virulence genes (4). One example is the *pil1* locus, which encodes the Pil1 pilus. This proteinaceous appendage at the bacterial surface allows *SGG* binding to collagen, bacterial colonization of cardiac valves, and the subsequent development of endocarditis (5).

Of medical interest, *SGG* was the first intestinal bacterium associated with colorectal cancer (CRC) (6). It is now widely accepted that colonic microbiota dysbiosis contributes to oncogenesis, with a higher relative abundance of potentially pro-carcinogenic bacteria, including *Escherichia coli pks*+, toxinogenic *Bacteriodes fragilis, Fusobacterium nucleatum*, *Parvimonas micra,* and *SGG* (7, 8). The *pks* island is a cluster of genes that encode polyketide synthases and nonribosomal peptide synthetases involved in the production of colibactin, which is a genotoxin that can cause DNA damage in eukaryotic cells. This toxin has been linked to increased mutation rates and has been implicated in the development of colorectal cancer (9).

A few studies have experimentally addressed the role of *SGG* in tumorigenesis (10–12). Our research group has shown that the *SGG* UCN34 strain colonizes the murine colon a 1,000-fold more effectively in a tumoral environment by outcompeting a phylogenetically-close member of the resident murine microbiota (11). More recently, we showed that UCN34 can also accelerate colon tumor development in a murine model by activating several cancer-related signaling pathways in epithelial and stromal colonic cells (12).

The oncogenic role of *E. coli* pks+ and enterotoxinogenic *B. fragilis* in colorectal cancer have been well documented (13). Specific genetic loci encoding the oncotoxin colibactin (*clbB, pks+*) and fragylisin also known as *B. fragilis* toxin (*bft*) have been identified within pathogenicity islands (14, 15). To determine whether unique genetic loci or pathogenicity islands are uniquely present in CRC-associated *SGG*, we performed de novo sequencing of five novel *SGG* isolates associated with CRC and compared them to 30 other isolates linked to various clinical conditions. Our research shows that *Streptococcus gallolyticus* subsp. *gallolyticus* (*SGG*) isolates linked to colorectal cancer (CRC) are not clonal but rather diverse. We did not identify any genetic cluster specific to the *SGG*-associated with CRC suggesting that the association between *SGG* and CRC is likely multifactorial and largely dependent on the host. Nevertherless, the presence of *SGG* could serve as a potential diagnostic marker for CRC.

## Materials and Methods

### Bacterial strains

Thirty-five *SGG* clinical strains isolated from blood samples were obtained from the National Reference Center for Streptococci in France (**Table 1**). They were grown at 37°C without shaking in Todd-Hewitt broth for genomic DNA isolation.

**Table 1.** Characteristics of whole-genome sequences of *Streptococcus gallolyticus* subsp. *gallolyticus*.

| Strain | Diagnostic | Size (bp) | Number of contigs | Number of coding sequences | G+C% | MLST type |
| --- | --- | --- | --- | --- | --- | --- |
| SGG17 | IE + polyps | 2,419,039 | 29 | 2,423 | 37.66 | 7 |
| SGG20 | IE + colon cancer | 2,300,175 | 25 | 2,295 | 37.49 | 91 |
| SGG21 | IE | 2,256,983 | 37 | 2,224 | 37.46 | 26 |
| SGG22 | IE | 2,392,006 | 27 | 2,366 | 37.81 | 15 |
| SGG23 | Non colon cancer | 2,421,052 | 31 | 2,429 | 37.66 | 7 |
| SGG24 | IE + Non colon cancer | 2,557,573 | 54 | 2,574 | 37.41 | 11 |
| SGG25 | IE + Non colon cancer | 2,429,569 | 38 | 2,471 | 37.49 | 7 |
| SGG26 | IE + Non colon cancer | 2,412,039 | 78 | 2,422 | 37.5 | 62 |
| SGG27 | IE + Non colon cancer | 2,401,851 | 30 | 2,416 | 37.49 | 11 |
| SGG28 | IE | 2,490,913 | 74 | 2,504 | 37.51 | 18 |
| SGG29 | IE | 2,300,216 | 23 | 2,292 | 37.49 | 91 |
| SGG30 | IE + colon cancer | 2,345,098 | 1 | 2,307 | 37.61 | 41 |
| SGG31 | NA | 2,310,255 | 19 | 2,315 | 37.59 | 5 |
| SGG32 | NA | 2,357,582 | 21 | 2,359 | 37.57 | 5 |
| SGG33 | IE + colon cancer | 2,317,420 | 29 | 2,322 | 37.43 | 5 |
| SGG34 | IE + colon cancer | 2,312,760 | 22 | 2,304 | 37.57 | 5 |
| SGG35 | IE | 2,239,166 | 26 | 2,23 | 37.54 | 5 |
| SGG36 | IE | 2,293,095 | 30 | 2,31 | 37.47 | 5 |
| SGG37 | IE + Colon cancer | 2,273,624 | 1 | 2,207 | 37.78 | 2 |
| SGG38 | IE + Polyps | 2,410,412 | 43 | 2,403 | 37.46 | 3 |
| SGG39 | Polyps | 2,667,828 | 149 | 2,764 | 37.27 | 118 |
| SGG40 | Spondylodiscitis + colon cancer | 2,468,312 | 2 | 2,492 | 37.67 | 5 |
| SGG41 | IE + colon cancer | 2,202,809 | 19 | 2,15 | 37.6 | 120 |
| SGG42 | IE + Colon cancer | 2,440,986 | 2 | 2,422 | 37.65 | 116 |
| SGG43 | IE + Polypes | 2,325,599 | 14 | 2,335 | 37.52 | 109 |
| SGG44 | IE + Colon cancer | 2,366,909 | 1 | 2,318 | 37.62 | 26 |
| SGG46 | IE + Diverticulosis | 2,445,476 | 43 | 2,487 | 37.46 | 121 |
| SGG47 | IE + Non colon cancer | 2,674,135 | 100 | 2,786 | 37.18 | 11 |
| SGG48 | IE | 2,357,783 | 21 | 2,36 | 37.49 | 7 |
| SGG49 | IE + Non colon cancer | 2,233,030 | 11 | 2,199 | 37.54 | 2 |
| SGG50 | IE + Non colon cancer | 2,310,202 | 43 | 2,329 | 37.52 | 43 |
| SGG51 | IE | 2,398,145 | 21 | 2,374 | 37.8 | 15 |
| SGG52 | IE + Non colon cancer | 2,375,034 | 64 | 2,407 | 37.47 | 119 |
| SGG53 | IE | 2,274,758 | 19 | 2,275 | 37.53 | 5 |
| SGG54 | NA | 2,304,878 | 16 | 2,287 | 37.56 | 2 |
| UCN34 | IE + Colon cancer | 2,350,911 | 1 | 2,307 | 37.64 | 41 |
| TX20005 | IE + Colon cancer | 2,258,003 | 1 | 2,224 | 37.73 | 117 |
| ATCC 43143 | NA | 2,362,241 | 1 | 2,246 | 37.52 | 12 |
| BAA 2069 | NA | 2,356,444 | 1 | 2,309 | 37.65 | 6 |
| DSM 16831 | Non human (Koala) | 2,492,900 | 1 | 2,464 | 37.7 | 1 |
| **Mean** |  | **2,372,680** |  | **2,368** | **37.55** |  |

IE, infective endocarditis; NA, Not available
Red, Cancer-associated strains; Green, No cancer-associated strains; Blue, Non human strain

### Antimicrobial susceptibility tests

Antimicrobial activity was tested by the agar disk-diffusion assay. A standardized inoculum of *SGG* strain was plated onto the surface of Mueller-Hinton agar containing 5% horse blood. Filter paper disk impregnated with antimicrobial agent are placed on the agar and plates were incubated overnight at 37°C. The isolates were tested for susceptibility to penicillin, ampicillin, tetracyclin, gentamicin, kanamycin, erythromycin, lincomycin, pristinamycin, rifampicin, vancomycin, teicoplanin, levofloxacin, linezolid, sulfamethoxazole/trimethoprim, norfloxacin and oxacillin. The minimum inhibitory concentration (MIC) of tetracyclin was determined using the E-test gradient strip method (Biomerieux, Marcy l’Etoile, France).

### Genomic DNA isolation, sequencing and assembly

Streptococcal strains were grown in Todd-Hewitt broth and incubated overnight at 37°C without agitation. The genomic DNA was extracted using DNAeasy blood and tissue kit (Qiagen) according to the manufacturer’s instruction.

Genome sequencing of five *SGG* strains associated to colon cancer (SGG30, SGG37, SGG40, SGG42, and SGG44) was performed in-house using a PacBio Sequel I instrument. Librairies were prepared according to the Pacbio Protocol for multiplex SMRT sequencing with v3 chemistry, utilizing the SMRTbell Express Template kit 2.0 and barcoded adapters kit. Assemblies were performed using the LORA pipeline v1.0.0 from Sequana (16). The LORA pipeline was run with default parameters and reproducible containers from the Damona project (https://github.com/cokelaer/damona), with Canu (17) as the assembler. Circularisation was performed (18) and coverage homogeneity was assessed using Sequana Coverage software for quality control. The final scaffolds consisted of a single circularized contig for SGG30, SGG37, and SGG44 and two contigs for SGG40 and SGG42. The resulting sequences were annotated with the RAST server (19).

Genome sequencing of additional 30 *SGG* strains was performed at the Singapore Centre for Environmental Life Sciences Engineering to at least a coverage of 100x on the Illumina Miseq platform (reagent kit v3, 2 x 300 bp). The paired-end sequencing reads were trimmed and assembled de novo into contigs on the CLC Genomics Workbench 10.0 (Qiagen). The resulting sequences were annotated with the RAST server (19). Raw sequencing reads used in this study are available on NCBI under accession number <u>PRJNA762634</u> (20).

### Comparative genomic analysis

REALPHY (reference sequence alignment based phylogeny builder) (21) web server was used to construct a phylogenetic tree using sequence data from our whole genome sequencing of 35 *SGG* and 1 *SGM* strains with default parameters. In addition, we used the complete genome sequence of five *SGG* strains, UCN34 (GenBank accession no FN597254) (3), TX20005 (GenBank accession no NZ_CP077423) (22), ATCC 43143 (GenBank accession no NC_017576) (23), ATCC BAA-2069 (GenBank accession no NC_015215) (24), and DSM16831 (GenBank accession no NZ_CP018822) (25) and two *SGM* strains ACA-DC198 (GenBank accession no HE613569) (26) and 679 (GenBank accession no GCA_900094105) (27) available in the NCBI database.

Jspecies web server (28) was used to determine the genetic diversity among the genomes by calculating the average nucleotide identity (ANI) based on ANIm (MUMer) and the OrthoVenn3 web server (https://orthovenn3.bioinfotoolkits.net/) was used to identify orthologous gene clusters in the genomes (default parameters: E-value 1×10-5, inflation value 1.5).

The core- and pan-genome were defined using SPINE/AGEnt/ClustAGE web server (29). MLST analysis based on on the divergence of sequence of seven housekeeping genes (*aroE, glgB*, *nifS*, *p20*, *tkt*, *trpD*, and *uvrA*) was performed in silico using the scheme developed for *SGG* by Dumke et al. (30). The MLST type of *SGG* strains was determined using the public databases for molecular typing and microbial genome diversity (pubMLST.org) (31).

Lipoprotein signal peptides and genes putatively involved in virulence were predicted by using SignalP-6.0 (32) and the virulence factor database (VFDB, https://www.mgc.ac.cn/cgi-bin/VFs/v5/main.cgi) (33), respectively.

Antibiotic-resistance genes were identified with the Resistance Gene identifier (RGI) program from the comprehensive antibiotic-resistance database (CARD) (https://card.mcmaster.ca/analyze/rgi) (34) and the putative genomic islands by using the IslandViewer 4 database (http://www.pathogenomics.sfu.ca/islandviewer/) (35).

## Results and Discussion

### Generalities and comparative genomics

De novo sequencing of five clinical strains of *Streptococcus gallolyticus* subsp. *gallolyticus* (*SGG*) using PacBio technology revealed an unexpected genetic diversity. These strains isolated from the blood of patients with colon cancer and designated SGG30, SGG37, SGG40, SGG42, and SGG44 were analyzed alongside two previously sequenced *SGG* strains associated with colorectal cancer, UCN34 (Accession number, NC_013798) and TX20005 (Accession number, NZ_CP077423). The general characteristics of these seven *SGG* strains are summarized in **Table 1**. The circular chromosome size ranged from 2,258,003 bp to 2,468,312 bp, with 2,207 to 2,492 open reading frames (ORFs). The G+C content remained consistent at 37.7%, across all seven strains. Each strain contained 18 predicted rRNAs while the number of predicted tRNA ranged from 70 to 72. Annotation using the RAST server indicated that approximately 23% of coding sequences (CDS) were classified as “hypothetical proteins”.

Additionally, whole-genome short-read sequencing of 30 clinical *SGG* strains recovered from human blood samples was performed by Illumina technology and de novo assembly. The number of contigs ranged from 11 to 149, while the estimated circular chromosome size varied between 2,202,809 to 2,674,135 bp (**Table 1**, **Fig. S1A**). The number of CDS ranged from 2150 to 2786 (mean, 2,368) (**Fig. S1B**). The average G+C% was 37.5% (37.2% - 37.8%) (**Fig. S1C**). Among the 30 additional *SGG* isolates of our collection, four (SGG20, SGG33, SGG34, and SGG41) were isolated from patients with colon cancer, while nine (SGG23, SGG24, SGG25, SGG26, SGG27, SGG47, SGG49, SGG50, and SGG52) were from patients without colon cancer (**Table 1**).

For further analysis, three additional genomes from the NCBI database were included: ATCC 43143, ATCC BAA 2069, and DSM 16831 (**Table 1**). Notably, DSM 16831 is the only non-human *SGG* strain, isolated from Koala feces. A whole-genome circular comparative map was generated for cancer-associated isolates (UCN34, TX20005, SGG30, SGG37, SGG40, SGG42, SGG44, SGG20, SGG33, SGG34, SGG41) colored in red and non-cancer associated isolates (SGG23, SGG24, SGG25, SGG26, SGG27, SGG47, SGG49, SGG50, SGG52) colored in green with DSM 16831 colored in blue also included for reference (**Fig. 1**). In this figure, genomes are organized according to their degree of identity (average nucleotide identity). This representation clearly shows the intertwining of red and green genomes indicating that *SGG*-isolates associated with CRC do not segregate from *SGG* belonging to the non-cancer group.

**Fig. 1.**
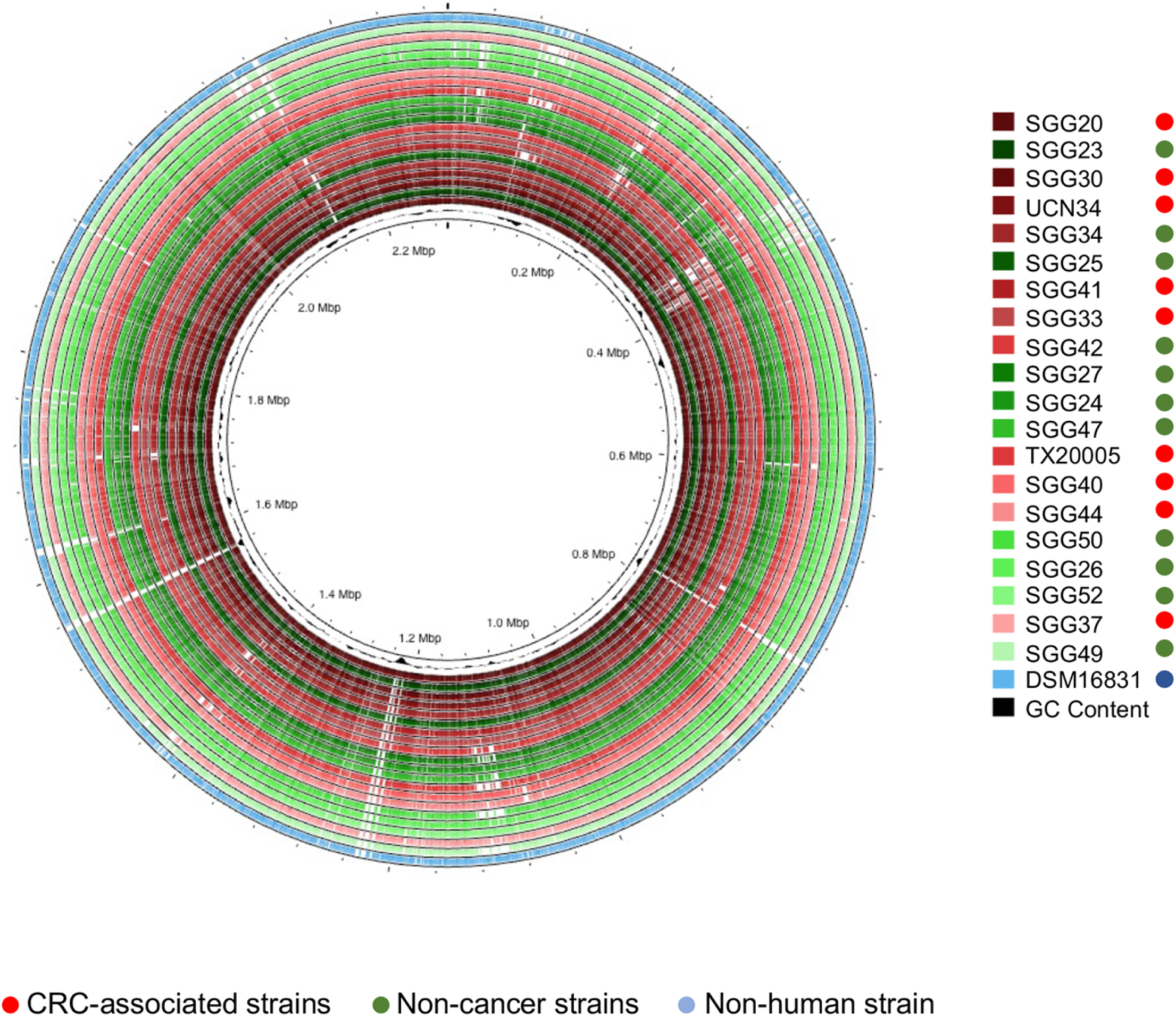
Circular representation of *SGG* genomes associated with colorectal cancer- (in red) or non cancer-associated (in green), including a non-human *SGG* isolate DSM16831 (in blue) for comparison. The inner-most circle (in black) shows the G+C% content plot. The genomes were classified based on their level of nucleotide identity.

Average nucleotide identity (ANI) analysis revealed that within the cancer-associated group, the SGG37 and SGG40 were the most genetically distant strains (ANI, 98.97%), while UCN34 and SGG30 were the most closely related (ANI, 99.99%) (**Table S1**). In the non-cancer group, SGG47 and SGG49 were the most distant (ANI, 98.93%), whereas SGG27 and SGG47 were the closest (ANI, 99.99%). The average ANI within the cancer and non-cancer group was 99.41% and 99.24%, respectively. Comparisons with the non-human strain DSM 16831 showed an average ANI of 99.03% with the cancer-associated group and 98.98% with the non-cancer group (**Table S1**).

### Phylogenetic trees

To assess the genetic diversity of *SGG* isolates, Multi Locus Sequence Typing (MLST), as established by Dumke et al (30) was employed analyzing seven housekeeping genes (*aroE*, *glgB*, *nifS*, *p20*, *tkt*, *trpD* and *uvrA*). The number of different alleles identified for each gene was as follows: *aroE* (8), *glgB* (8), *nifS* (7), *p20* (10), *tkt* (7), *trpD* (9) and *uvrA* (10). In total, 22 Sequence Types (ST) were detected among the 40 *SGG* strains, indicating high genetic diversity (**Table 1**). The most common STs were ST5 (8/40, 20%), ST7 (4/40, 10%), ST 2 (3/40, 7.5%), ST11 (3/40, 7.5%), ST41 (3/40, 7.5%), and ST91 (3/40, 7.5%). Additionally, six novel STs were identified: ST116 (SGG42), ST117 (TX20005), ST118 (SGG39), ST119 (SGG52), ST120 (SGG41), and ST121 (SGG46). Fourteen STs were unique (14/40, 35%). No correlation was observed between MLST types and cancer association. Among the 11 *SGG* isolates from cancer patients, eight distinct STs were represented, while the 10 strains from non-cancer patients were distributed across seven STs, respectively (**Table 1**).

A high-resolution phylogenetic tree was generated using the whole genome sequences of the 40 *SGG* and 3 *SGM* strains (**Fig. 2**). The distribution of strains revealed that those isolated from blood of patients with tumors were not clonal but highly diverse.

**Fig. 2.**
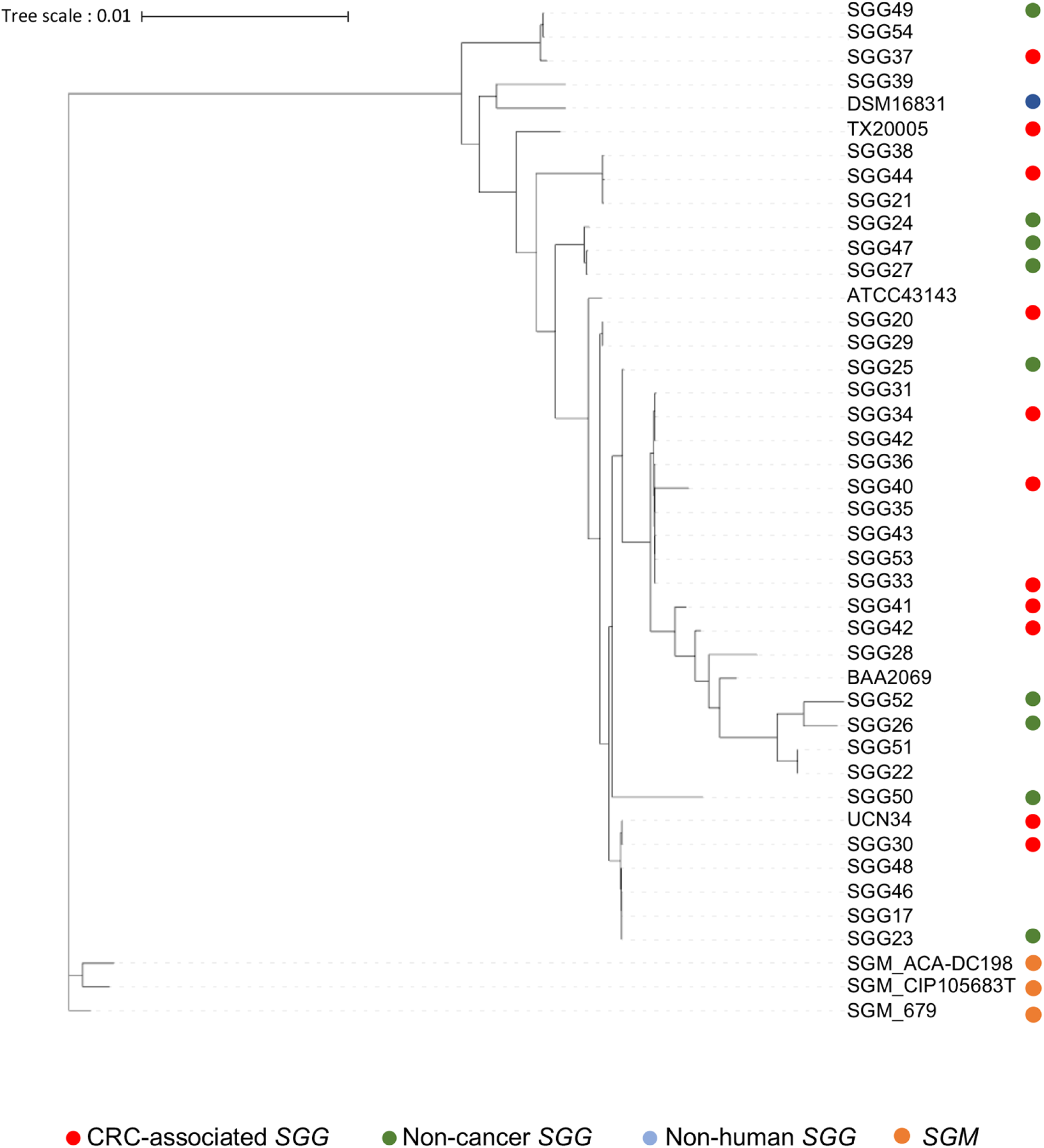
Phylogenetic tree based on whole genome sequence data of 40 *S. gallolyticus* subsp. *gallolyticus* and 3 *S. gallolyticus* subsp. *macedonicus* strains. The tree was constructed using REALPHY (21) and managed with iTOL web site (52).

### Determination of the core- and pan-genomes

The core genome consists of sequences conserved across all strains, while the pan-genome comprises the core, the accessory genome (genes present in at least 2 genomes), and strain- or species-specific genes (found in only one genome). The core- and pan-genomes of *SGG* strains from both cancer- and non-cancer groups were determined using the “Spine and Agent” web server (http://spineagent.fsm.northwestern.edu/index_age.html) (29). The total number of predicted coding sequences (CDS) and the average GC content for each group are summarized in **Table 2**. The core genome size for strains associated with cancer (n=11) and those not associated with cancer (n=9) was 1.64 Mbp and 1.63 Mbp, respectively, accounting for an average of 80.1% and 75.2% of the total genome (**Table 2**).

**Table 2.** *SGG* core- and pan-genome characteristics.

|  | Cancer- associated group (n=11) |  | Non cancer-associated group (n=9) |  |
| --- | --- | --- | --- | --- |
|  | Core | Pangenome | Core | Pangenome |
| Size (nucleotides) | 1,646,264 | 2,950,941 | 1,629,376 | 3,181,519 |
| % of genome | 80.1 (76.2-84) |  | 75.2 (68.1-81.1) |  |
| G+C % | 38.8 | 37.5 | 38.8 | 37.4 |
| Nb pred. CDS | 1,856 | 3,727 | 1,833 | 4,137 |
| Gene length (bp) | 887 | 792 | 889 | 769 |

The size of the pan-genome depends on the number of strains analyzed. For the cancer-associated and non-cancer-associated groups, the pan-genome sizes were 2.95 Mbp and 3.18 Mbp, respectively (**Table 2**).

A comparison of the entire genomes of *SGG* isolates from cancer and non-cancer groups did not reveal any specific genes or genetic locus (**Fig. 3**). Unlike well-characterized colorectal cancer (CRC)-associated bacteria, such as *Escherichia coli* pks+ and *Bacteroides fragilis* bft+, which encode the oncotoxins colibactin and fragylisin, respectively, *SGG* genome analysis did not identify any genetic island potentially encoding an oncotoxin directly linked to cancer development.

**Fig. 3.**
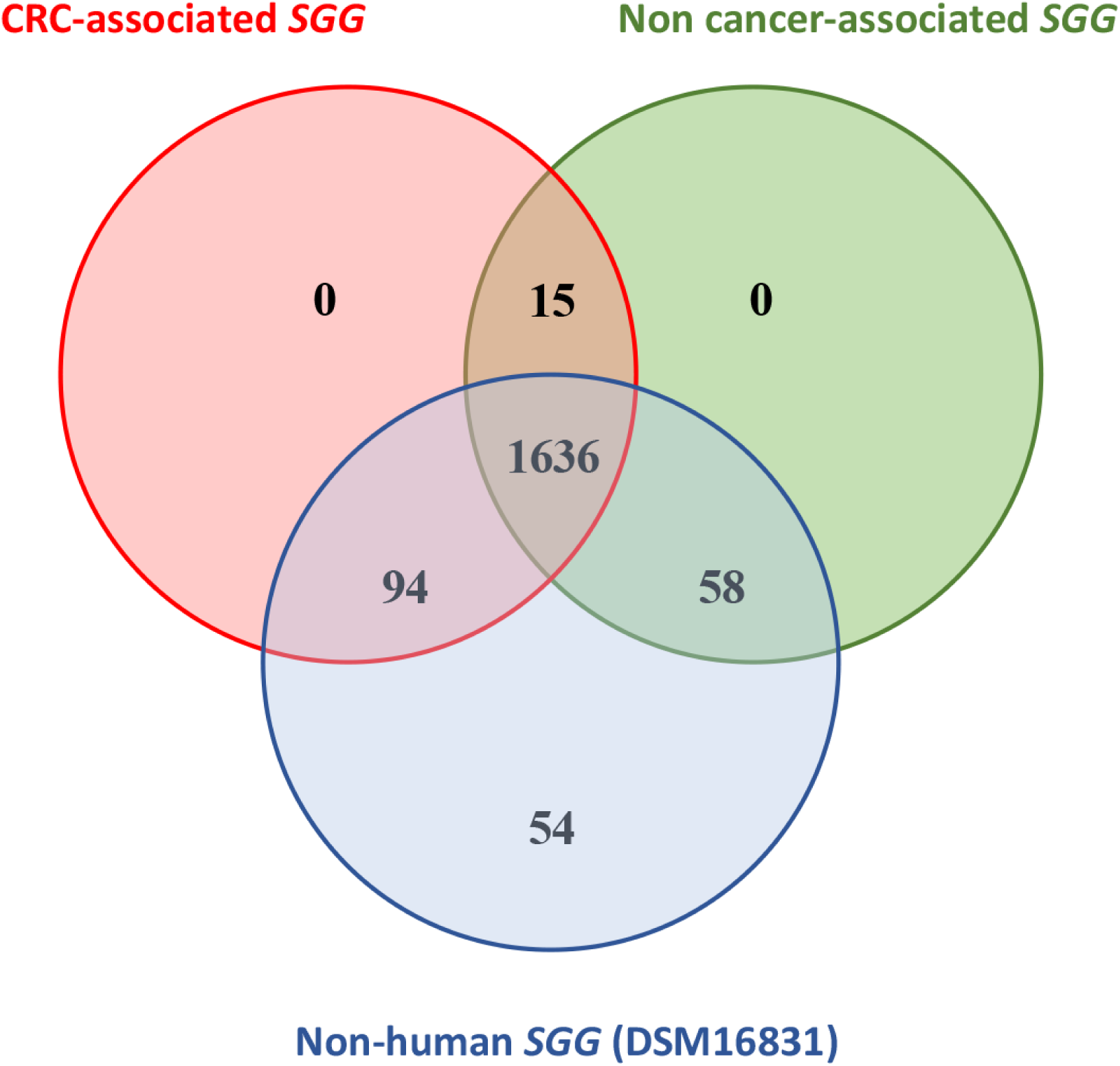
Venn diagram representing common and specific orthologous clusters of *SGG* core genomes from cancer associated- , non cancer associated-, and a non human- strains. The numbers of unique and shared orthologous clusters of each core genome is indicated.

### Genomic islands

Horizontal gene transfer (HGT) enables the acquisition of new gene in both prokaryotic and eukaryotic genomes. Genomic islands (GI) are regions acquired through HGT that may harbor genes implicated in metabolism, pathogenicity or antibiotic resistance.

Putative genomic islands were identified using the IslandViewer4 webtool (35). Among the *SGG* strains associated with cancer, the number of GIs ranged from 4 to 15, with GI-to-genome size ratios spanning from 4.5% to 11.6% (**Table 3**).

**Table 3.** Determination of genomic islands in cancer-associated *SGG* (red), non cancer-associated *SGG (green)*, non-human *SGG* (blue) and in S. gallolyticus subps. *macedonicus* (orange)

|  | Cancer-associated SGG |  |  |  |  |  |  |  |  |  |  |  | Non cancer-associated SGG |  |  |  |  |  |  |  |  |  | Non human SGG | SGM |  |  |  |
| --- | --- | --- | --- | --- | --- | --- | --- | --- | --- | --- | --- | --- | --- | --- | --- | --- | --- | --- | --- | --- | --- | --- | --- | --- | --- | --- | --- |
|  | UCN34 | TX20005 | SGG20 | SGG30 | SGG37 | SGG40 | SGG42 | SGG44 | SGG33 | SGG34 | SGG41 | Mean | SGG23 | SGG24 | SGG25 | SGG26 | SGG27 | SGG47 | SGG49 | SGG50 | SGG52 | Mean | DSM16831 | ACA_DC198 | 679 | CIP_105683T | Mean |
| No of GIs | 8 | 13 | 7 | 8 | 11 | 15 | 8 | 9 | 9 | 10 | 4 | 9.9 | 8 | 15 | 9 | 12 | 9 | 15 | 5 | 9 | 14 | 11.0 | 9 | 16 | 15 | 16 | 15.7 |
| GI total length (kb) | 171 | 186 | 129 | 171 | 185 | 287 | 174 | 243 | 169 | 160 | 98 | 193 | 148 | 265 | 265 | 279 | 217 | 361 | 86 | 231 | 290 | 238 | 169 | 333 | 261 | 390 | 328 |
| Largest GI (kb) | 36 | 41 | 39 | 36 | 40 | 43 | 53 | 64 | 47 | 46 | 46 | 44 | 36 | 47 | 72 | 47 | 62 | 97 | 38 | 43 | 47 | 54 | 47 | 95 | 94 | 66 | 85 |
| Mean length (kb) | 21 | 14 | 18 | 21 | 17 | 18 | 22 | 27 | 19 | 16 | 25 | 20 | 18 | 18 | 29 | 23 | 24 | 24 | 17 | 43 | 21 | 24 | 19 | 21 | 17 | 24 | 21 |
| Size GI/Size genome (%) | 7.4 | 8.2 | 5.6 | 7.4 | 8.6 | 11.6 | 7.1 | 10.6 | 7.3 | 6.9 | 4.5 | 8.4 | 6.1 | 10.4 | 10.9 | 11.6 | 9.0 | 13.5 | 3.8 | 8.5 | 12.2 | 9.8 | 7.3 | 15.6 | 12.4 | 17.6 | 15.2 |

Genomic islands were determined by using Island viewer 4 (http://www.pathogenomics.sfu.ca/islandviewer/) (35).

### Antibiotic resistance genes

Analysis of the 40 *SGG* genomes using the CARD database revealed that at least one antibiotic resistance gene was present in 85% of the strains (**Table S2**). Tetracyclin resistance was highly prevalent in *SGG* strains, with the genes *tet(M)* and *tet(O)*, which confer tetracycline resistance through ribosome protection, and *tet(L)*, which encodes a tetracycline efflux protein, detected in 28/40 (70%), 7/40 (17.5%) and 4/40 (10%) strains, respectively. Overall, 82.5% of *SGG* strains carried at least one gene that could confer resistance to tetracycline. Tetracycline, a broad-spectrum antibiotic, is widely used in the livestock industry and in veterinary treatment. In 2005, Leclercq et al. showed that tetracycline resistance was widespread in *SGG* with a frequency of tetracycline resistance of 77.7%, similar to that observed in our study (36). Furthermore, they also found that *tet(M)* was the most prevalent determinant of resistance to tetracycline, since it was present in 97% of tetracycline-resistant isolates, similar to our *SGG* group where it was present in 87% of TetR isolates. As suggested for *Streptococcus agalactiae* (37), in which the dominance of *tet(M)* was also noted, resistance to tetracycline is commonly observed in *SGG* likely due to the extensive use of tetracycline.

A gene cluster (*aadE*-*sat4*-*aphA-3*), originally described as part of the transposon *Tn*5405 in methicillin-resistant *Staphylococcus aureus* (38) was identified in 17/40 *SGG* strains. This cluster confers resistance to streptomycin (*aadE*), streptothricin (*sat4*), and kanamycin (*aphA-3*), and was often found alongside the *erm(B)* gene, which confers resistance to macrolides, lincosamides and streptogramins B. Additionally, a *dfrF* gene, involved in trimethoprim resistance, was detected in one strain (SGG28), and a *catA9* gene, which confers resistance to chloramphenicol, was found in two strains (SGG33 and SGG52) (**Table S2**). Resistance to streptomycin, erythromycin, or chloramphenicol was confirmed in strains harboring *aadE, erm(B)* or *cat* gene, respectively (data not shown).

Antimicrobial resistance testing of all *SGG* strains using disk diffusion assay confirmed that the antibiotic resistance genes detected in silico were expressed in vitro and that their repertoires could confer different levels of resistance (data not shown). For example, two genes implicated in tetracyclin resistance, *tet(M)* and *tet45*, were found in UCN34, while only one gene, *tet (M)*, was found in TX20005. In vitro tetracycline resistance determination using E-test strips clearly showed that UCN34 was more resistant to tetracyclin than TX20005, with measured MIC values of 48 µg/ml and 16 µg/ml, respectively. Representative antibiograms performed in routine for Streptococci are shown in **Fig. 4** for the six sequenced *SGG* strains (SGG20, 30, 37, 40, 42, 44), the reference strains UCN34 and TX20005, and *SGM* CIP105683T for comparison. SGG40 and SGG44 were resistant to multiple antibiotics whereas TX20005 and SGG37 were amongst the most sensitive strains. Like many other Streptococci, most *SGG* isolates were found resistant to norfloxacin and oxacillin.

**Fig. 4.**
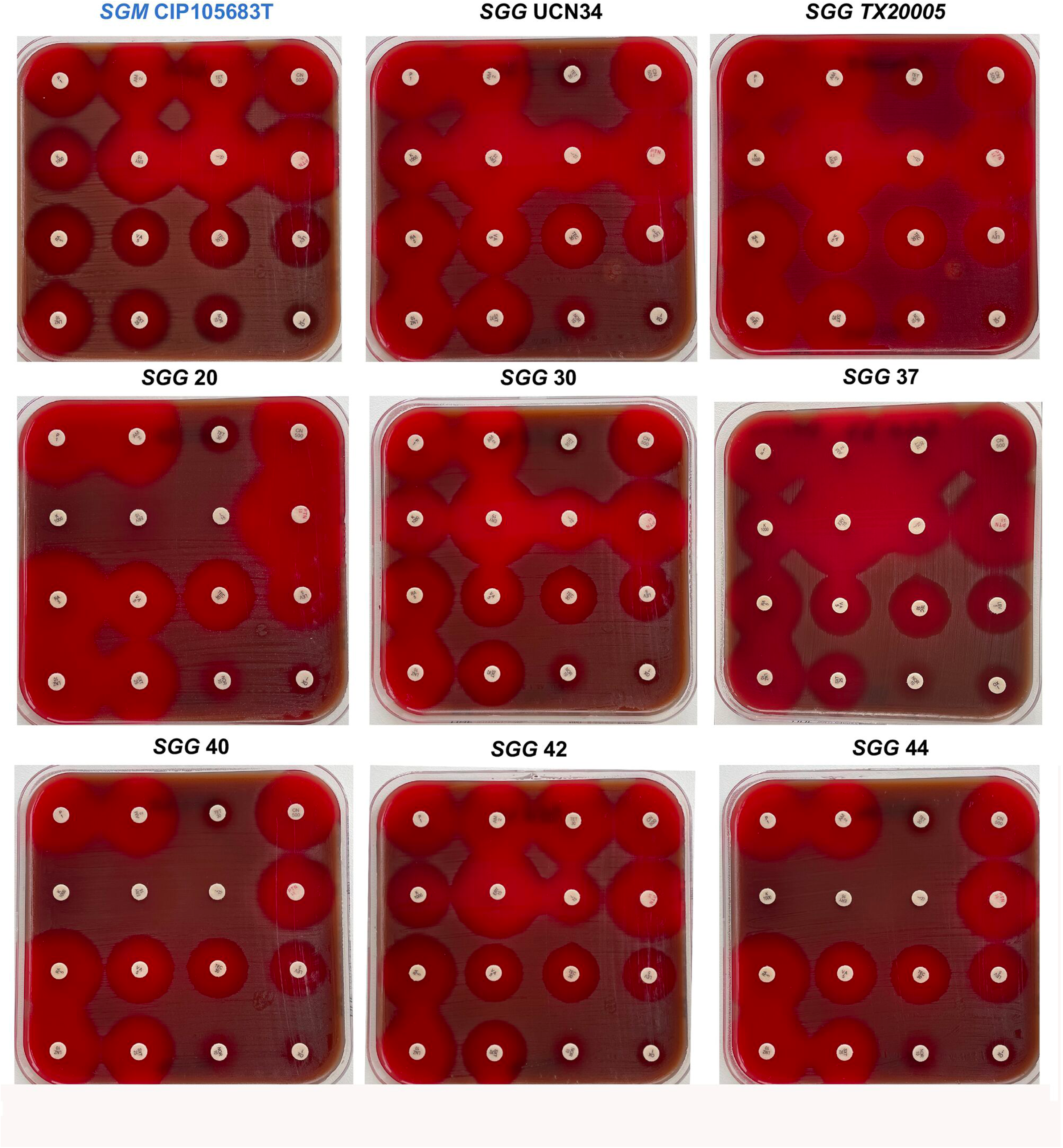
Visual representation of the antimicrobial susceptibility profile of six novel *SGG* isolates associated with colon cancer (SGG20, SGG30, SGG37, SGG40, SGG42, SGG44). As controls we included the reference *SGG* strains UCN34 and TX20005 and the non-pathogenic SGM CIP105683T. Each disc is labeled with the abbreviation of the antibiotic tested with the dose indicated below (in µg) Abbreviations: PEN, penicillin; AMP, ampicillin, TET, tetracyclin; CN, gentamicin; K, kanamycin; ERY, erythromycin; L, lincomycin; PTN, pristinamycin; RA, rifampicin; VA, vancomycin; TEC, teicoplanin; LEV, levofloxacin; LNZ, linezolid; SXT, sulfamethoxazole/trimethoprim; NOR, norfloxacin: OX, oxacillin.

### Predicted surface proteins

Surface proteins play a crucial role in bacterial invasion, adherence, and interactions with the host immune system and environment. These proteins can be categorized into three groups based on their mode of attachment: LPxTG proteins (anchored at C-terminal end), hydrophobically bound proteins, and lipoproteins (anchored at the N-terminal end) (39).

#### LPxTG proteins

LPxTG proteins are located on the bacterial surface and anchored to the cell wall by sortase A (40, 41). An analysis of *SGG* strains from both cancer and non-cancer groups identified a diverse repertoire of proteins containing both an N-terminal signal peptide and a C-terminal LPxTG motif. Most of these proteins are predicted to be involved in adherence to host proteins (**Table 3**). Of the 18 LPxTG proteins detected in the UCN34 strain (3), only six (33%) were present in all genomes of cancer-associated strains (**Table 4**). Among these, five were also found in non-cancer strains.

**Table 4.**
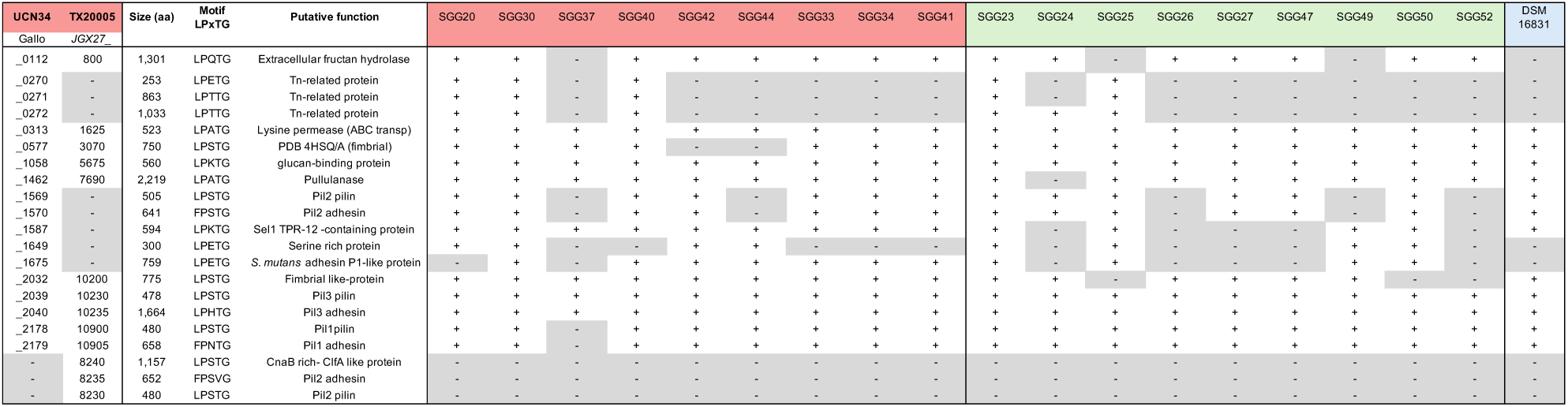
List of LPxTG proteins of UCN34 and TX20005 present or absent in cancer associated - group (red), non cancer- associated group (green) and non human isolate (blue)

Sortase A, a transpeptidase enzyme, anchors covalently LPxTG proteins to bacterial cell-wall. In the UCN34 genome, three genes encoding sortase A were identified: the housekeeping Gallo-1127 (present in all 40 *SGG* strains), Gallo_0299 (found in 20/40 strains, 50%) and Gallo_1651 (present in 22/40 strains, 55%). These genes, designated as *srtA*, *srtA_1_*, and *srtA_2_*, are located downstream of genes encoding GyrA, plasmid mobilization proteins, and a “phage tail tip lysozyme”, respectively. Additional sortase genes, *srtA*_3_ (SGG24, SGG42), *srtA*_4_ (SGG47) and *srtA*_5_ (SGG40) were sporadically detected with the number of *srtA* genes varying from 1 to 4 among *SGG* isolates. All *srtA* genes encode potentially functional enzymes with the characteristic catalytic motif TL(V/I)TC at the C-terminal region, a hallmark of the SrtA protein (42). Analysis of the genomic context of these *srtA* genes (**Fig. S2**) revealed that (i) the *srtA* gene, present in all strains, is consistently downstream of *gyrA*; (ii) additional *srtA* genes (*srtA_1_-srtA_5_*) are consistently found near mobile genetic elements (e.g. transposase, integrase), suggesting that while *srtA* serves as the primary housekeeping enzyme, auxiliary *srtA* genes were acquired through horizontal gene transfer. Blast analysis confirmed that homologs of these additional sortases are found in other *Streptococcus* species, such as *SGM*, *S. gallolyticus* subsp. *pasteurianus* (*SGP)*, *Streptococcus infantarius*, and *Streptococcus lutetiensis*.

#### Pili

In Gram-positive bacteria, covalently-assembled pili play a key role in adherence and host tissue colonization (43). The UCN34 genome contains three loci involved in pilus biosynthesis: Pil1 (Gallo_2177-2179), Pil2 (Gallo_1570-1568), and Pil3 (Gallo_2038-2040) (3). Pil1 and Pil3 have been extensively studied in UCN34. Pil1 is essential for adhesion to collagen type I and contributes to heart valve colonization and endocarditis in a rat model (5), while Pil3 facilitates adhesion to colonic mucus and colonization of the distal colon in mice (44). Both *pil1* and *pil3* were detected in all 40 *SGG* strains. The *pil1* cluster, located between a *tetR*-like gene, encoding a transcriptional regulator and a trehalose catabolism locus was conserved in 33 strains. In the remaining 7 strains, two additional genes encoding a DNA repair protein and a signal peptidase were inserted upstream of the *pil1* locus (**Fig. S3**). In these strains, the genes coding for the accessory and major pilins were more distantly related to those in UCN34. Notably, in the CRC-associated SGG37, the genes encoding the major pilin and sortase C in the *pil1* locus were absent. The Pil1 locus is replaced by a large SpaA isopeptide-forming pilin related protein of 1,571 amino acids. (**Fig. S3**). The *pil3* gene cluster was highly conserved in all *SGG* strains, with at least 90% identity across the locus. However, three genes, downstream of *pil3*, encoding a copper transport system, a protease, and a lactate dehydrogenase, were absent in five strains (SGG37, SGG39, SGG49, SGG54 and the non-human isolate DSM 16831) compared to the other 35 other *SGG* strains. Unlike *pil1* and *pil3*, the *pil2* gene cluster was less conserved and was absent in 10/40 (25%) of *SGG* strains. Of note, *pil2* was absent in two CRC-associated isolates (SGG37 and SGG44). When present, it was well conserved, except in TX20005 (**Fig. S3**). Interestingly, a specific *pil2* locus was detected in the genome of TX20005 and was not found in any of the other 38 strains studied. Overall, these results highlight the critical role of Pil3, which contributes to mucus-binding and host colon colonization by *SGG*. Pil1 locus, involved in collagen binding and endocarditis, is also well conserved except in a few isolates. Pil2 appears as a species-specific locus whose role remains to be determined.

#### Lipoproteins

Lipoproteins are anchored to the bacterial membrane via an N-terminal lipid moiety attached to a conserved cysteine residue. They are involved in various biological processes, particularly in transport and sugar uptake systems. Compared to LPxTG proteins, lipoproteins exhibit lower variability within *SGG* strains. Using the SignalP 6.0 web server, the number of predicted lipoproteins in *SGG* genomes ranged from 39 to 47. Of the 43 putative lipoproteins identified in UCN34, 34 (79%) of them were conserved across all cancer-associated isolates (**Table S3**). One lipoprotein, Gallo_1778, was found in all cancer-associated strains except strain TX20005 and was mostly absent (7 out of 9) in non cancer-associated isolates and the non-human isolate DSM 16831. This protein contains a CarG-like motif, which is associated with carbapenem intrinsic resistance factor, and a glycosyl hydrolase motif, suggesting a potential role in bacterial survival and host interactions.

### Biosynthesis of various extracellular polysaccharides in *SGG* strains

Extracellular polysaccharides play a crucial role in bacterial ability to evade the immune response and to survive in hostile environments. In *SGG* UCN34, three distinct genetic loci have been identified as contributing to polysaccharide biosynthesis.

#### Capsule

A cluster of 12 genes (*cpsA-M*) enables *SGG* UCN34 to synthesize an extracellular capsule (3). Like other extracellular pathogens, this capsule protects *SGG* from phagocytosis and clearance by macrophages (45). Previous studies have shown that the *cps* locus, which encodes the biosynthesis of capsular polysaccharides biosynthesis in *SGG*, consists of two parts: a highly conserved 5’ region and a species- or strain-specific 3’ region (46). The present analysis of 40 *SGG* strains supports this observation. The first 5 genes (*cpsA-E*), located in the 5’ region of the locus, are highly conserved, while the 3’ region varies among strains, containing between six and eleven genes. Based on nucleotide sequence analysis, we identified nine distinct *cps* cluster types. The capsule type of each *SGG* isolate is indicated in **Fig. 5**. Our results confirms that CRC-associated *SGG* are diverse and belong to 6 different capsular types.

**Fig. 5.**
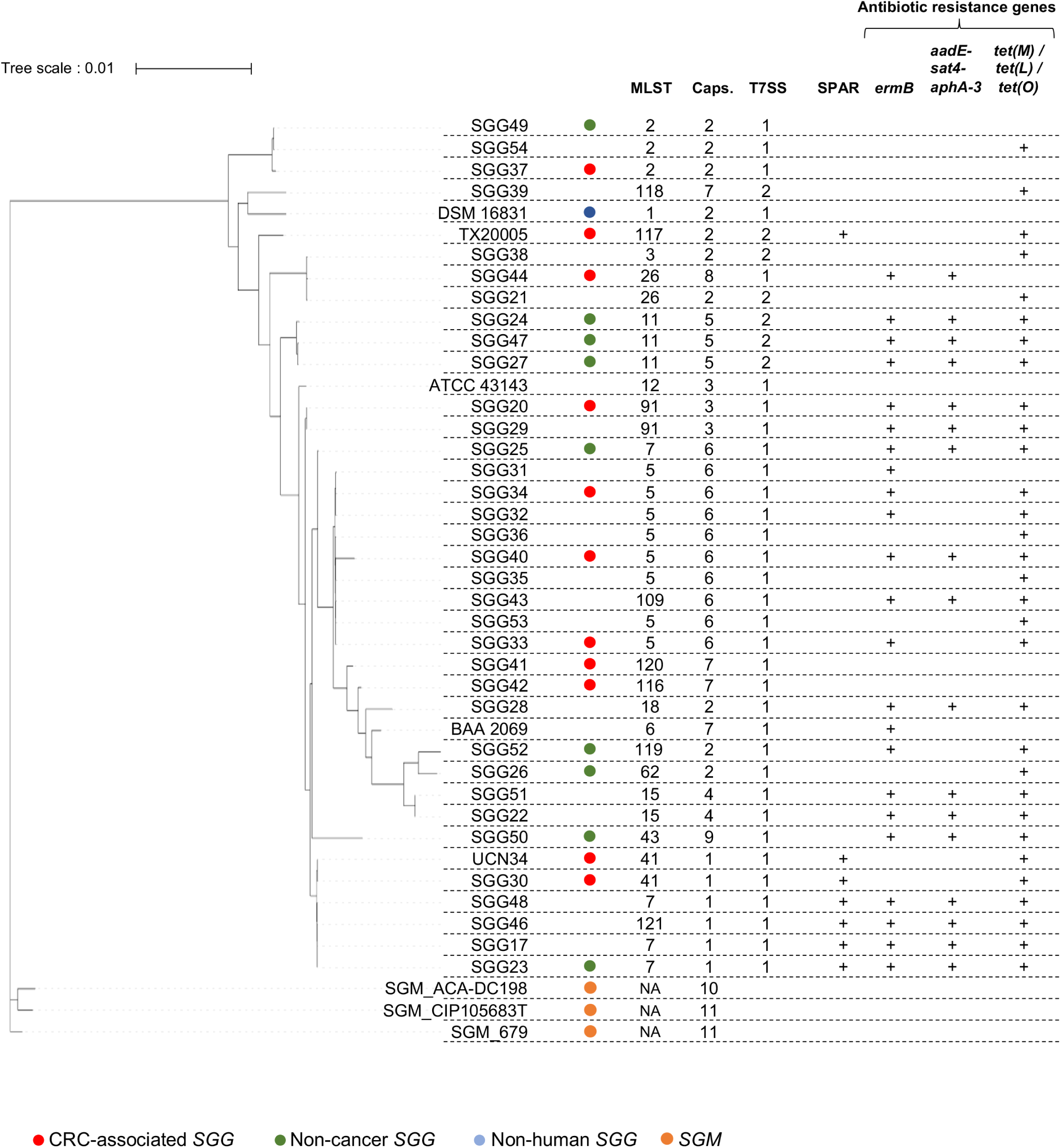
Graphical abstract of this study. Phylogenetic tree based on whole genome sequence data of 40 *S. gallolyticus* subsp. *gallolyticus* and 3 *S. gallolyticus* subsp. *macedonicus* strains. The tree was constructed using the REALPHY webserver (21) and managed with iTOL web site (52). Type of T7SS was arbitrarily defined as 1 for T7SS_UCN34_-type and 2 for T7SS_TX20005_-type.

#### Hemicellulose

A genetic locus (*gallo_0364-0367*) putatively involved in hemicellulose biosynthesis has been identified in UCN34 (3). This locus consists of four genes encoding a putative diguanylate cyclase, two glycosyltransferases, and a hypothetical transmembrane protein. Analysis of our *SGG* genome collection revealed that this locus is present and highly conserved across the 40 *SGG* strains. However, a few strains exhibit single nucleotide polymorphisms (SNPs) and stop mutations, likely leading to locus inactivation (**Fig. S4**).

A similar locus is present in *S. infantarius*, *S. lutetiensis*, *S. equinus*, *S. salivarius*, and *S. ruminicola* but is absent in *Streptocococcus gallolyticus* subsp. *pasteurianus* (*SGP).* In *SGM* CIP105683T, this locus is present, though the two genes encoding glycosyltransferases appear nonfunctional due to premature stop codons, and the fourth gene encoding the hypothetical transmembrane protein is missing.

#### Glucan

A glucan biosynthesis locus (*gallo_1052-1057*) encoding three glycosyl transferases and associated regulatory proteins has been described in UCN34 (3). These transferases are highly similar to *Streptococcus mutans* GtfA, GtfB and GtfC, which play key roles in bacterial adherence to tooth surfaces and biofilm formation (3).

Analysis of the 40 *SGG* genomes revealed that this locus is present in all strains. In most isolates (62.5%), the genetic organization is identical to that of UCN34 (**Fig. S5**). However, variations were observed in some strains: four strains (SGG20, SGG29, SGG42, and ATCC43143; 10%) lacked the first regulatory gene. Ten strains (25%) were missing both the first glycosyltransferase gene and its regulatory gene. Two strains (SGG31 and SGG36) contained a premature stop codon, leading to a truncated glycosyltransferase. One strain (SGG24) lacked the two last genes of the cluster (**Fig. S5**).

### Type VII secretion system

A locus encoding a type VII secretion system (T7SS) has been identified in *SGG* strains TX20005 (22) and UCN34 (20). In the TX20005 strain, the T7SS appears to play a role in colon colonization and the development of colon tumors (22).

While the 5’ region of the T7SS locus, which encodes the structural components of the secretion apparatus, is relatively conserved between the two strains, the gene cluster located downstream of *essC,* which encode putative effectors, shows significant differences (20).

Analysis of 38 additional *SGG* genomes revealed that T7SS associated with UCN34 (T7SS_UCN34_-type) is more commonly found than the T7SS associated with TX20005 (T7SS_TX20005_- type) (32 vs 8 strains, 80% vs 20%). Among the colorectal cancer (CRC) group, 9 out of 11 strains harbor the T7SS_UCN34_-type, compared to 7 out of 10 in the non-cancer group. The results are summarized in **Fig. 5**.

### TX20005 pathogenicity locus (*sparA-L*)

A recent study identified a chromosomal locus in *SGG* cancer-associated strain TX20005, designated the *SGG* pathogenicity-associated region (SPAR), which consists of twelve genes (JGX27_RS05685 to JGX_RS05740). This locus has been implicated in the adherence of strain TX20005 to colonic cells and its ability to stimulate CRC cell proliferation (47). It was observed that the Δ*spar* mutant phenocopies the Δ*esx* mutant (T7SS knock-out) in TX20005. It was proposed that this could be due to the presence of a putative transcriptional activator (JGX27_RS05710) of the T7SS system in the SPAR locus. An analysis of the 40 *SGG* genomes revealed that the entire locus was present in only 10 genomes (25%). Specifically, it was detected in 3 out of 11 (27.2%) *SGG* isolates associated with cancer and in just 1 out of 11 (9.1%) isolates from the non-cancer group (**Fig. 5**). These findings suggest that the SPAR locus may have a more limited role in tumor development than previously thought (47). Very interestingly, the only gene of the SPAR locus conserved in all 40 *SGG* is the putative transcriptional activator (JGX27_RS05710) described above. These results demonstrate the importance of studying several clinical strains and the role of National Reference Center in providing such contemporary circulating strains.

### Virulence factors

The presence of putative virulence genes in the *SGG* genomes was investigated using the VFDB database. Genes associated with adherence, including fibronectin-binding protein (*fbp*), glucosyltransferases (*gtf*), sortases (*srt*), plasmin receptor protein (*plr*) or lipoprotein rotamase A (*slrA*), were identified in *SGG* (**Table S4**). Additionally, a few proteases, such as C3-degrading protease, C5a peptidase, and HtrA serine protease were detected, which may help *SGG* to evade the host immune system. The prevalence of these potential virulence factors is summarized in **Table S4**.

Bile salt hydrolase (BSH), also known as choloylglycine hydrolase, encoded by *bsh* gene, is commonly found in many bacterial species, especially those present in the gastrointestinal microbiota, such as *Lactobacillus*, *Bifidobacteria*, *Clostridia*, and *Enterococci*. The *bsh* gene is strictly conserved in all *SGG* strains. The potential role of these virulence and resistance factors in the pathogenicity of *SGG* warrants further investigation, particularly in the context of host-pathogen interactions.

### Comparison S. gallolyticus subps. gallolyticus vs S. gallolyticus subsp. macedonicus

*S. gallolyticus subsp. macedonicus* (*SGM*) is a thermophilic, homofermentative, dairy *Streptococcus*, that is phylogenetically very close to *SGG*. Unlike *SGG* and *SGP*, *SGM* is considered as non-pathogenic (48), which is reflected by a reduced number of pathogenicity-related genes compared to *SGG* (4). Previous sequencing of *SGM* CIP 105683T (49), a type strain from the Institut Pasteur Collection, was used to perform detailed genomic analysis including two others publicly available *SGM* genomes: ACA-DC 198 (Accession number, NC_016749) and 679 (Accession number, GCA_900094105). The general features of these strains are presented in **Table 5**.

**Table 5.** Characteristics of whole-genome sequences of three *S. gallolyticus subsp. macedonicus* (*SGM*)

| **Strain** | **Acc. Number** | **Size (bp)** | **G+C content** | **ORFs** | **Coding %** |
| --- | --- | --- | --- | --- | --- |
| ACA-DC198 | NC_016749 | 2,130,034 | 37.6 | 2,248 | 86.2 |
| 679 | GCA_900094105 | 2,107,416 | 37.5 | 2,277 | 86.3 |
| CIP_105683T | PRJNA940176 | 2,210,410 | 37.7 | 2,372 | 85.7 |

The core genome of *SGM* subspecies was determined from these three strains. Next, a comparison of *SGM* and *SGG* core genomes revealed that 117 proteins, belonging to 30 different orthologous clusters, were specific to the *SGM* core genome (**Fig. S6**). Our findings support the data reported by Papadimitriou at al (4) regarding the absence of several virulence factors in *SGM* compared to *SGG* (**Table S4**).

*SGM* ACA-DC 198 produces two food-grade lantibiotics, macedocin and macedovicin. Macedocin is putatively encoded by two genes (SMA_1380 and SMA_1381) (50) within a 10-gene cluster, whereas macedovicin is encoded by a single gene (SMA_1409) (51). The gene cluster responsible for macedocin biosynthesis was also detected in *SGM* 679 and *SGM* CIP 105683T, but the macedovicin gene was found only in ACA-DC 198. In sharp contrast, all 40 *SGG* isolates harbor the genetic cluster encoding the class IIb bacteriocin named gallocin A which enables *SGG* to outcompete *Enterococcus faecalis*, a resident member of the host microbiota in tumoral conditions (11).

A large LPxTG protein of 7,961 amino acids (SMA_1990 in the ACA-DC 198 genome), was identified in *SGM*_ACA-DC 198. This protein contains a YSIRK signal peptide motif at the N-terminus, five Flg-new (Pfam 09479) domains, and 48 cadherin repeat-like (cI46864; Pfam 05345) domains at the C-terminus. The Flg-new domain, found in intestinal archea such as *Methanomassiliicoccales*, exhibits structural similarity with mucus binding domains. Interestingly, the Pil3 adhesin (Gallo_2040) which mediates mucus binding, also contains four Flg-new domains (**Fig. S7**). The same LPxTG protein was found in *SGM* CIP 105683T, but with only 28 cadherin repeat-like domain, whereas it was absent in SGM_679. SMA_1990 homologs were identified in various *Streptococcus* species, including *S. infantarius, S. thermophilus, S. iniae*, *S. parauberis*, *S. suis*, *S. dysgalactiae*, as well as in *Lactococcus* species such as *L. lactis* and *L. raffinolactis*. However, it was absent in the two subsp. *SGG* and *SGP*.

Unlike *SGG*, *SGM* lacked the gene clusters encoding the type VII secretion system (T7SS) and glucan biosynthesis. The absence of T7SS, a system often associated with bacterial ability to compete with closely related bacteria and colonize the gut microbiota, may further explain the non-pathogenic nature of *SGM*. Similarly, the lack of glucan biosynthesis genes, which contribute to biofilm formation and adhesion in pathogenic *Streptococcus* species, suggests that *SGM* has a reduced ability to establish persistent infections. Moreover the gene encoding bile salt hydrolase is probably non functional in *SGM* due to a premature stop codon in the coding sequence. These genetic differences highlight the evolutionary divergence between *SGM* and its pathogenic counterpart, *SGG*, reinforcing the hypothesis that *SGM* is adapted for a non-pathogenic, dairy-associated lifestyle rather than a virulent, host-associated one.

### Concluding Remarks and Perspectives

The analysis of 11 clinical *Streptococcus gallolyticus subsp. gallolyticus (SGG)* genomes from patients with colorectal cancer (CRC) and their comparison with isolates from non-CRC patients did not identify a specific subtype of strains associated with disease. One key finding of this study is the high diversity among CRC-associated *SGG* isolates, which belong to various MLST types, exhibit different capsule compositions, and contain diverse T7SSb loci (**Fig. 5**). Furthermore, the TX20005 pathogenicity locus (SPAR), previously described as a potential cancer -associated determinant (47), is not conserved in *SGG*, and was detected in only three out of eleven CRC-associated strains. However, the putative regulator of the Type VII secretion system detected in the SPAR locus, is present in all the SGG sequenced so far. Notably, the majority of human-derived *SGG* isolates (87%) carried at least one gene conferring resistance to an antibiotic while the non-human strain, DSM 16831, lacked any antibiotic resistance genes. This suggest that antibiotic resistance in human *SGG* isolates were likely acquired in vivo through horizontal gene transfer (HGT). These findings highlight the potential impact of host environment on the genetic adaptation of *SGG*. In sum, these genomic analyses highlight the opportunistic nature of *SGG* and its complex association with cancer relying mostly on *SGG*’s ability to colonize the host colon and its ability to compete with bacterial members of the host colon microbiota through production of specific bacteriocin such as gallocin A and other LXG-effectors secreted by the specialized T7SSb whose role need to be further studied.

## Acknowledgements

We are grateful to Nicolas Dmytruk for his help in the retrieving of *SGG* isolates from CNR and Chloé Plissonneau for her expertise in antibiograms. We are thankful to Christophe Rusniok et Laura Gomez-Valero for discussion and sharing their expertise in bioinformatic analyses. We thank Philippe Glaser and Asmaa Tazi for critical reading of the manuscript.

This work was supported by the National Research Foundation and Ministry of Education Singapore under its Research Centre of Excellence Program (SCELSE). S. Dramsi acknowledges the support of the Institut National contre le Cancer (INCA; grant PLBIO16-025) and from the French Government’s Investissement d’Avenir program, Laboratoire d’Excellence Integrative Biology of Emerging Infectious Diseases (grant no. ANR-10-LABX-62-IBEID).

## Supplementary figures legend

**Fig. S1**. Whole genome characteristics of 40 *S. gallolyticus* subsp. *gallolyticus* strains.

**Fig. S2**. Schematic representation of the genetic localisation of the various *srtA* genes.

**Fig. S3**. Schematic representation of the various loci involved in pili biosynthesis. Arrows indicate sense of transcription. Nomenclature indicated in the upper locus is accordingly to UCN34 (3).

**Fig. S4**. Hemicellulose biosynthesis. Schematic representation of the locus involved in hemicellulose biosynthesis in the 40 *S. gallolyticus* strains. Arrows indicate sense of transcription. Nomenclature indicated in the upper locus is accordingly to UCN34.

**Fig. S5**. Glucan biosynthesis. Schematic representation of the locus involved in glucan biosynthesis in the 40 *S. gallolyticus* strains. Grey and dashed grey arrows indicate genes encoding glycosyltransferases and regulators, respectively. Nomenclature indicated in the upper locus is accordingly to UCN34.

**Fig. S6**. Venn diagram showing the distribution of shared orthologous clusters among core genomes of CRC-, non CRC-associated strains, and SGM. The numbers of unique and shared orthologous clusters of each core genome is indicated.

**Fig. S7**. Schematic representation of predicted domains within SMA_1190 of SGM_ACA-DC198 (upper) and Gallo_2040 protein of UCN34 (lower) using the Simple Modular Architecture Research Tool web server (SMART, http://smart.embl-heidelberg.de/).

